# Attention Guided Mechanism Interpretable Drug-Gene Interaction (MIDI) Modeling for Cancer Drug Response Prediction and Target Effect Explanation

**DOI:** 10.1101/2025.03.31.646490

**Authors:** Tingyi Wanyan, Ling Cai, Deepak Nijhawan, Bruce Posner, Jiwoong Kim, Rongqing Yuan, Ruheng Wang, Jingwei Huang, Shengjie Yang, Prema Mallipeddi, Tao Wang, Xiaowei Zhan, Guanghua Xiao, Yang Xie

## Abstract

Cancer drug discovery using genetic information is still poorly developed. Precisely locating drug atoms and explaining the targeting effect is crucial in precision medicine since it helps understand the drug’s mechanism of action. Much data has been collected regarding drug response against cancer cell lines, and many models predict the drug response based on genomic information. However, to our knowledge, none of the data-driven techniques propose to detect the targeting mechanism of small drug molecules against genetic targets. In this work, we propose MIDI (Mechanism Interpretable Drug-Gene Interaction) model to delve deep into the targeting relation between drug molecules against genetic patterns. We show that purely based on a data-driven approach, the attention mechanism in our model could capture the important binding effect of small molecules towards gene targets. We provide both theoretical derivation and experiment results to show the information flow regarding the attention mechanism. In the meantime, we demonstrate that our model presents much higher prediction performance with the interpretation mechanism than the other state-of-the-art drug response prediction models.

## Introduction

Precision medicine in cancer drug discovery aims to tailor the most effective therapy to individual patients. Due to the high heterogeneities in different cancer patients’ genetic screenings, chemotherapy with old-designed drug structures typically shows much less effectiveness on diverse cancer types with increasing tumor drug resistance. Newly effective cancer drug discovery becomes a more urgent request with the rapid growth of global cancer patient populations, approximately 2 million new patients each year^1^. Meanwhile, identifying new, effective drugs for cancer patients is of great importance to improve survival rates and reduce treatment-related toxicities. Traditional approaches to cancer drug discovery heavily depend on the experience of manually selecting the possible drug structures, with the hope that some may work on certain cancer cell lines. However, drug screening on particular drug structures is both time-consuming and resource-consuming, the experiment cycles and clinical trials typically take months or even years to complete, which counts for billions of dollars in investment. The failing rate is high, approximately 90% of the drug candidates fail due to the inaccuracy of the pre-clinical drug structure design^2^; in other words, the assumption of a newly designed unknown drug structure targeting specific gene mutations is eventually proven incorrect. Therefore, AI-aided precise medicine on cancer drug discovery in the pre-clinical phase becomes a much more attractive direction. To date, even though substantial resources have been invested in generating vast amounts of data to establish the relationship between molecular characteristics of cells and drug response in complex cancer types^3-5^, AI data-driven computational approaches remain poorly developed in understanding the drug MOA(mechanism of action) on targeting specific mutated genes, as well as facing the significant challenges exist in predicting cancer drug responses.

More specifically, traditional computational methods such as linear regression-based^6-8^, kernel-based^9-11^, and ensemble-based^12,13^ models have struggled with high-dimensional data, such as the vast quantity of genes, and have difficulty capturing the complex, non-linear connections between drug response and molecular characteristics. This leads to only modest prediction performance and fails to adequately explain the mechanisms behind cancer drug-gene targeting. In contrast, recent advances in deep learning have shown potential for enhancing prediction accuracy. However, many of these deep learning architectures utilize black-box feed-forward layers, offering little to no insight into the binding mechanisms of drug chemical structures or the interaction mechanisms with gene targets. Additionally, this limitation hinders the ability of these models to generalize and make predictions for new, unknown drug structures. Recently proposed protein-ligand targeting models. Many recently proposed DDI models apply self-attention graph structures to interpret drug compound properties, but these models use data purely from drug structures, with no connection between genetic targets. Thus, they can’t explain the drug-gene target effect,

In this work, we propose an original transformer-like model whose architecture is specifically designed to tackle the problems mentioned above, the model is named MIDI (Mechanism Interpretable Drug-Gene Interaction) shown in Fig. 1. We illustrate that with our unique design on MIDI, we can decently address the three major problems in the precision medicine domain: Target identification; Precise drug response prediction; Drug MOA interpretation. The novel design of our model can be briefly summarized in the following aspects: 1. Design unique model architectures to process drug molecular structure and its gene target effect, adopt the state-of-the-art foundation model Geneformer to extract the gene identification embeddings; 2. Design a detailed drug graph structure to make precise drug MOA explanation; 3. Design efficient drug-gene cross-attention architecture to process the vast amount of gene input; 4. Design unique loss function to incorporate drug-gene targeting interaction prior knowledge precisely; 5. Build a generalizability model architecture for training a potential drug-gene interaction foundation model efficiently and easily. With the above model architecture design, firstly, we demonstrate that MIDI shows mass increments over other baseline models on drug effect prediction tasks, on average 30% increment over the base line models; Secondly, MIDI is the first AI model designed to process the drug-gene target effect based on our knowledge, and we show that by capturing this targeting effect through the unique drug-gene cross-attention modeling, MIDI could precisely cluster the drug categories, as well as precisely predict the drug response on the cancer cell lines with mutated targeted genes; Thirdly, with the unique detailed design on the drug Graphformer structure, we demonstrate that MIDI could identify the most important atoms that perform the hydradon bond to the target protein residual, and detect the precise binding site on the drug chemical structure, this process is crucial in cancer drug discovery since it helps identify the leading small molecular component in designing drug candidate for a certain type of cancer, as well as helps understand the drug MOA. We test the model’s abilities through vast, carefully designed experiments to support our claims.

**Fig. 1:**
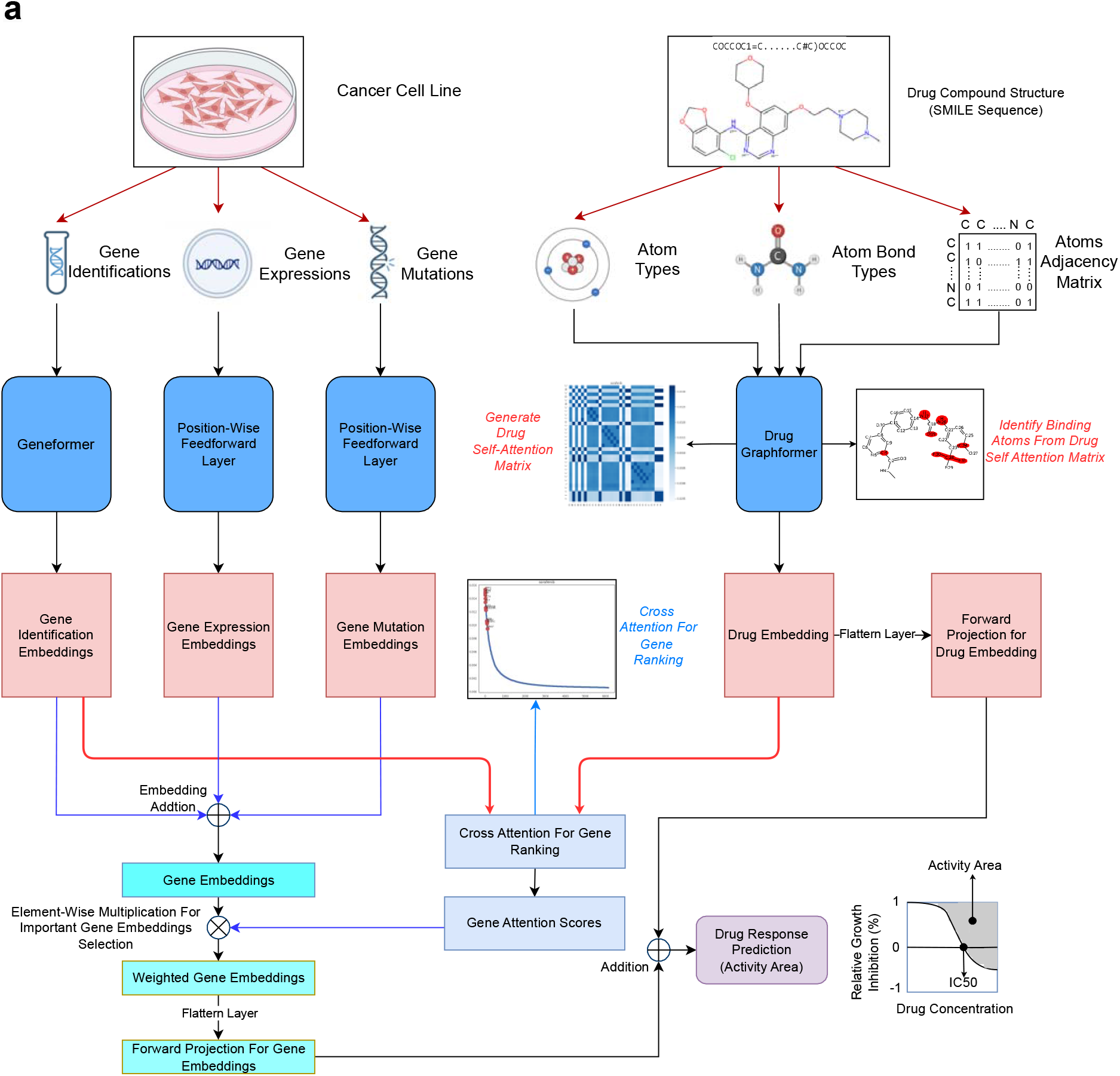
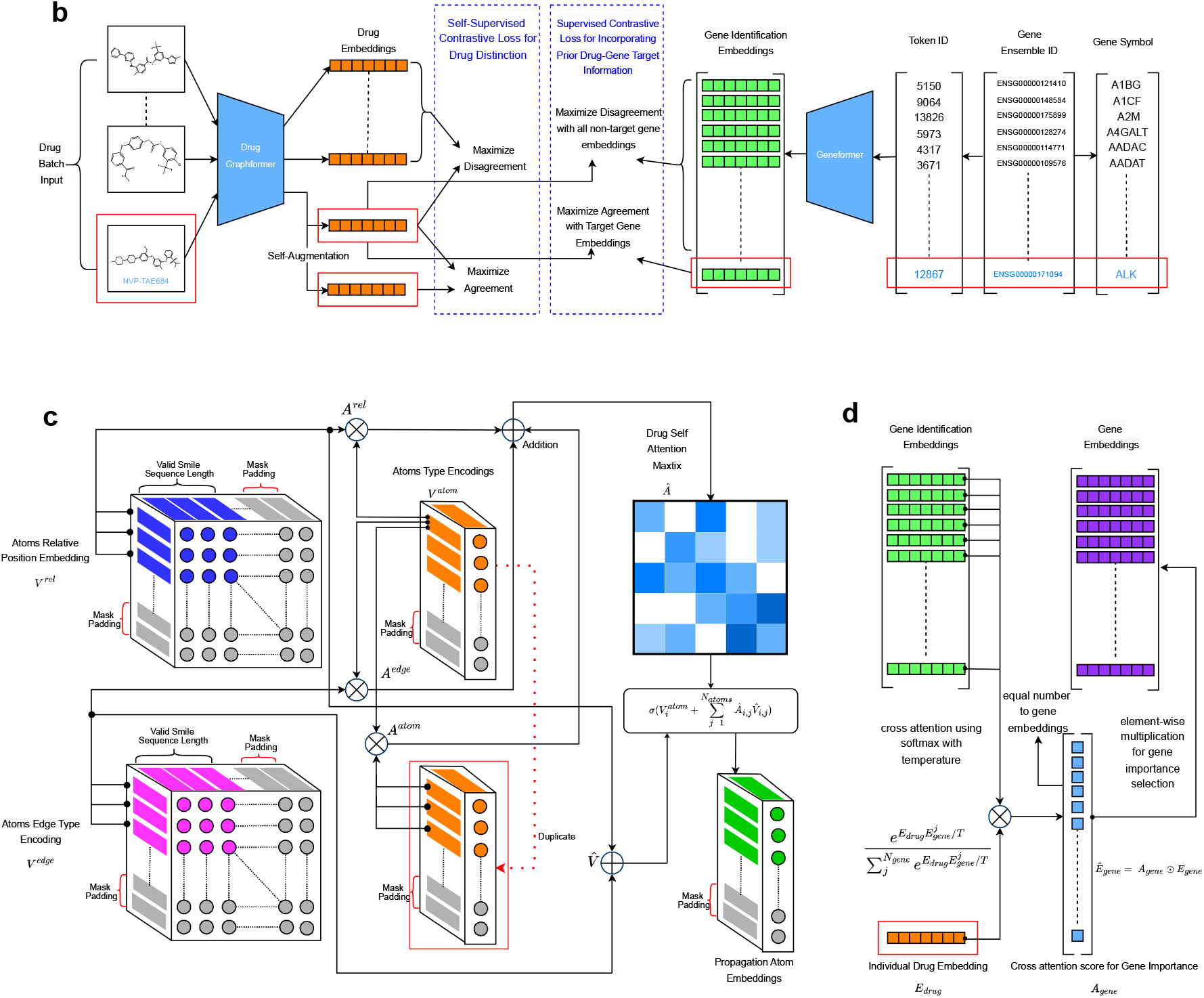
Detailed Technology Pipeline, Model Architecture, and Loss Functions. **a**. Technology pipeline and general model architecture. **b**. Learning strategy of incorporating prior knowledge on drug-gene interaction. **c**. Detailed illustration of the self-attention GraphFormer architecture for processing drug chemical structure input. **d**. Detailed illustration for Drug-Gene cross-attention architecture design for important gene ranking and selection.

Finally, the drug target effect explanation is crucial in cancer drug discovery, and we believe that through the unique architecture design, MIDI poses great potential for application in the massive pre-clinical drug screening phase to help extract the most crucial and valuable drug chemical component, this provides great help to narrow down the drug candidates for a given genomic profile with realistic pharmacogenetic bases. In the meantime, the unique design of the drug Graphformer combined with the global drug-gene cross-attention architecture makes it easy to concurrently train on millions of Drug-Cancer cell line pairs to construct a foundation model, which possesses the knowledge of drug-gene interaction information through drug chemical structure, as well as the knowledge to explain the drug MOA, these potential advantages make MIDI efficiently and easily to generalize the prediction and explanation on new, unseen drugs.

## Experiments and Results

### Data Description and Data Pre-Process

We use the famous publicly available data CCLE(Cancer Cell Line Encyclopedia)^6^ for our experiment and model training. It contains nearly 1700 human cancer cell lines across 39 primary cancer types with comprehensive molecular fingerprints, including mutation, gene expression, copy number variation, miRNA expression, protein array, and DNA methylation. We collect a total of 471 cancer cell lines that contain drug screening together with 24 drugs. The specific drug names are shown in the x-axis notification of Fig. 2D, and Fig. 2B shows the cancer cell line proportion for different cancer types.

**Fig. 2:**
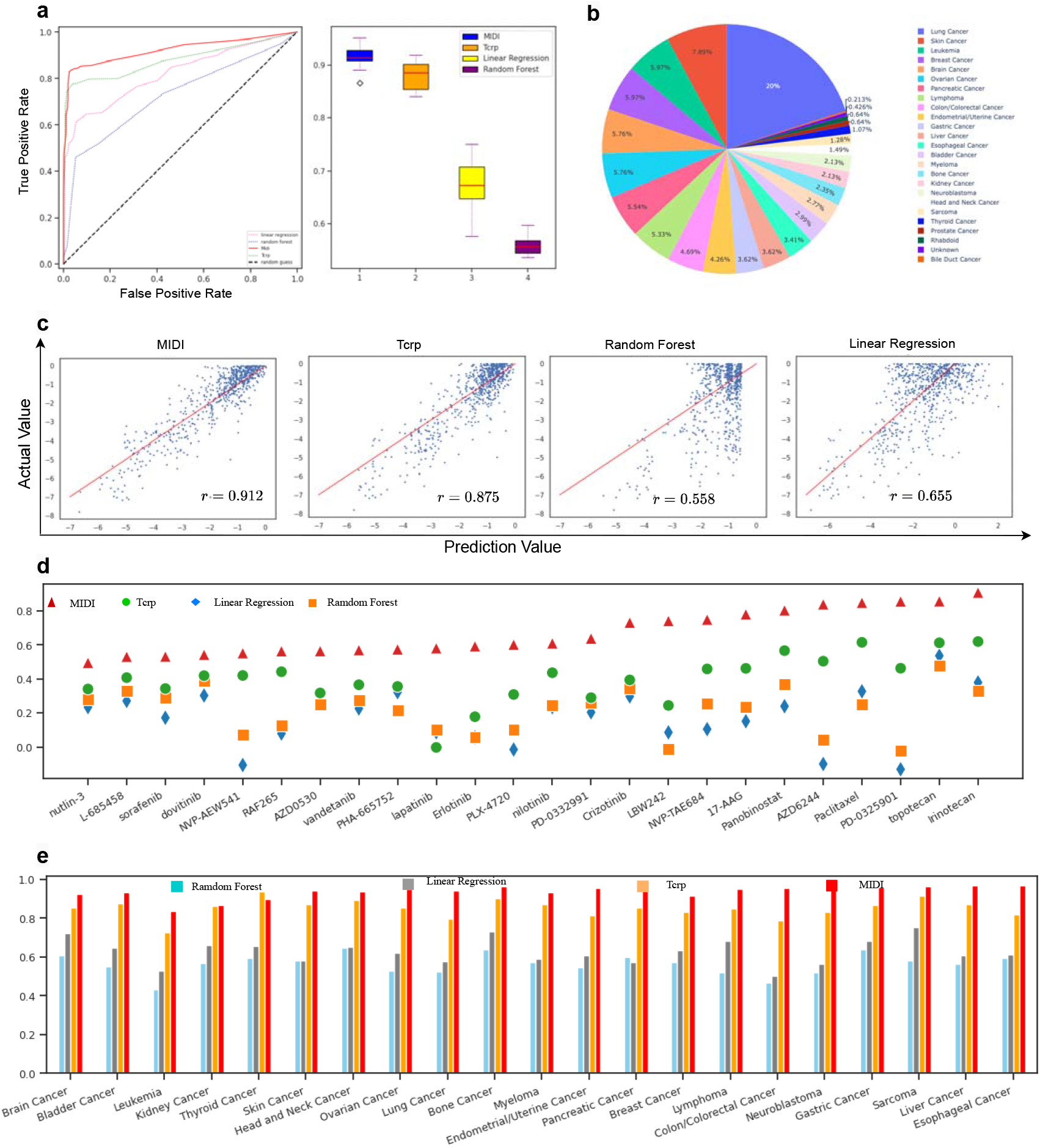
General Performance Comparison On Cancer Drug Effect Prediction. **a**. AUROC curve and 5-fold cross-validation for evaluating general performance on drug response prediction. **b**. The cancer type ratio for all cancer cell line data, total cancer cell line number is 471. **c**. Scatter plot for different model predictions. **d**. Per-Drug prediction comparison for different models, the y-axis is the correlation score, and the x-axis is different drug names. **e**. Per cancer type prediction comparison for different models, the y-axis is the correlation score, and the x-axis is the different cancer types.

To generate the comprehensive drug-gene interaction prior knowledge dataset for model training, we integrated data from multiple sources, including the Drug Gene Interaction Database (DGIdb), Genomics of Drug Sensitivity in Cancer (GDSC), and PubChem. Drug-gene interaction data was initially downloaded from DGIdb and processed to map drug names to PubChem Compound IDs. Canonical SMILES representations for compounds were retrieved using PubChem’s REST API. Drug-target interaction data from GDSC was processed to extract PubChem Compound IDs and target gene IDs. Interaction scores were derived from DGIdb annotations and aggregated across sources to ensure consistency and eliminate duplicates. Gene symbols and names associated with drug targets were mapped to NCBI Entrez Gene IDs using the gene_id.py script. This script parses the gene_info.gz file from NCBI FTP to extract mappings between gene symbols, synonyms, and Entrez Gene IDs, allowing taxonomy-specific filtering. For unresolved gene queries, external APIs such as NCBI E-utilities, HGNC, and Ensembl REST were used to retrieve Entrez Gene IDs. By standardizing gene identifiers across all datasets, this step ensured a consistent representation of genes in the final dataset. The resulting dataset integrates the following information: canonical SMILES, PubChem Compound ID, NCBI Entrez Gene ID, and interaction score.

For visualizing the drug, protein-ligand binding structure as shown in Fig. 5A for experiment Drug Molecule Binding Site Detection, we specifically conduct the following procedures: Firstly, compound identifiers were obtained from the PubChem Compound database using Entrez Programming Utilities (E-utilities) provided by NCBI (https://www.ncbi.nlm.nih.gov). Protein-ligand structural data were sourced from the Protein Data Bank (PDB, https://www.rcsb.org). Ligands, referred to as PDB heterogens, were mapped to their corresponding PDB structures using the Ligand Expo dataset (http://ligand-expo.rcsb.org/dictionaries/cc-to-pdb.tdd). To establish links between PDB heterogens and PubChem Compound identifiers, InChIKeys were extracted from the Chemical Component Dictionary (https://files.wwpdb.org/pub/pdb/data/monomers/components.cif) and queried against the PubChem database. Gene-to-PDB associations were derived from UniProt’s ID mapping datasets (https://www.uniprot.org). These mappings were integrated to associate ligand structures with their corresponding protein targets in PDB files. Protein-ligand interactions were analyzed using PyMOL (https://www.pymol.org). Hydrogen bonds were identified based on a donor-acceptor distance threshold of 3.3 Å, where donor atoms were defined as nitrogen or oxygen atoms bonded to hydrogens, and acceptor atoms as oxygen or nitrogen atoms not bonded to hydrogens. Hydrophobic interactions were identified between nonpolar carbon atoms in ligands and hydrophobic residues in proteins within a distance cutoff of 3.8 Å. Hydrophobic residues were defined as alanine (ALA), valine (VAL), leucine (LEU), isoleucine (ILE), methionine (MET), phenylalanine (PHE), tryptophan (TRP), and proline (PRO). Protein residues involved in these interactions were visualized in PyMOL. Hydrogen bonds and hydrophobic interactions were highlighted with distinct colors to facilitate interpretation.

### General Performance

We test MIDI’s drug effect prediction performance against traditional computation methods (Linear regression; Random Forest) and the state-of-the-art drug effect prediction model TCRP^14^. The data is divided into 80% for training and 20% for testing. Specifically, since the number of different cancer cell line types varies, as shown in Fig. 2B, the training and testing data is divided based on cancer cell lines as follows: For each type of cancer cell line, we pick 80% for training, and the remaining cell lines as testing data, repeat this procedure for all the cancer cell line types, then add the training data and testing data for different cancer type cell lines as the final training and testing data. There are certain cancer cell line types with very few cell line numbers, such as Bile Duct Cancer and Prostate Cancer, we put all cancer types with less than five cancer cell lines as the training data. Eventually, we had 376 cancer cell lines for training and 95 cancer cell lines for testing. Since the 24 drugs screened all the cancer cell lines, we have 9024 Drug-Cancer Cell Line pairs for training and 2280 for testing.

For all the baseline models (Linear Regression, Random Forest, and TCRP), the data input is the concatenation of the drug smile sequence and gene expression sequence. The loss function is the MSE loss for Linear Regression and Random Forrest, for TCRP, we adopt the same few-shot learning strategy as discussed in the original paper^14^. We first test the drug effect prediction correlation score on testing data. Fig. 2 A shows the AUC curve of the testing result on the left and the 5-fold cross-validation evaluation results on the right, MIDI shows the highest prediction performance, with an average of 5% increment over TCRP, 27% increment over Linear Regression, and 37% increment over Random Forest. Fig. 2C is the scatter plot that visually shows the prediction correlation performance. Next, we perform a per-drug effect prediction comparison. Specifically, we group the testing Drug-Cancer Cell Line pairs by different drug names and collect the prediction results for each drug. The prediction results are shown in Fig. 2D. MIDI shows a marginal performance increment on per-drug effect prediction over all other baseline models, on average 25% increment over TCRP, 40% over Linear Regression, and 45% over Random Forest, the reason for this marginal performance increment is specifically discussed in the Discussion Section. Finally, we perform the drug effect prediction comparison in a per-cancer-type manner. Specifically, the testing data is grouped into different cancer types, and the prediction results are collected accordingly. Fig. 2E shows the testing results. Again, MIDI shows the highest prediction performance against baseline models for all the cancer types except for Thyroid Cancer. The reason is that there are only five cancer cell lines for Thyroid Cancer, where we use three cell lines for training and two for testing. In such a scenario with very little training data, MIDI shows degraded performance compared to TCRP since TCRP adopts a few-shot transfer learning strategy that specifically deals with few training data. This suggests incorporating a few-shot learning strategy into our model training for possible future development.

### Gene Cross-Attention Score for Drug-Gene Targeting Effect Interpretation and Hierarchical Drug Clustering

To our knowledge, no other current AI models could incorporate the drug-gene interaction information nor explain the drug-gene target effect. In this experiment, we demonstrate our model’s ability to explain the Drug-Gene targeting effect through the unique gene cross-attention design after incorporating prior knowledge. To fully establish the ability to interpret, we use all the cancer cell line data as the training data. Specifically, we trained the model with 50 epochs on all the Drug-Cancer Cell Line pairs with the batch size set to 50. The detailed loss functions incorporating prior knowledge are discussed in Section Loss Function. After training, we specifically examine the cross-attention scores for the gene ranking part of the model, Fig. 3A shows the hierarchical clustering on the 24 drugs based on the distribution of the cross-attention scores for the whole 6144 genes, we discover that the scores correctly cluster most of the drug pairs with similar MOA which specifically listed as text in the middle of Fig. 3a, this shows the model after training has a strong ability to recognizes different categories of the drugs by their MOA through the corresponding gene cross-attention ranking scores. Note that each drug has a unique cross-attention gene ranking score since drug MOA is fixed when its chemical structure is fixed. We then specifically pick six pairs of drugs from the drug hierarchical clustering plot with similar MOAs and plot the gene cross-attention scores in a ranking order. Fig. 3B shows the gene ranking attention scores for six drug pairs, and Fig. 3c shows the top 20 genes with the highest attention scores. Firstly, the gene ranking plots show that the most important targeted genes for specific drugs are all ranking very high, for example, EGFR and SRC rank top one for Erlotinib and AZD0530. This shows the model correctly identifies the targeted genes for a specific drug with the incorporated prior knowledge through specifically designed supervised contrastive loss. Interestingly, the number of targeted genes for a specific drug structure is typically very few. However, given the prior knowledge of the few targeted genes, the model could correctly cluster the drug types based on the attention scores over 6144 genes through the data-driven training strategy.

**Fig. 3:**
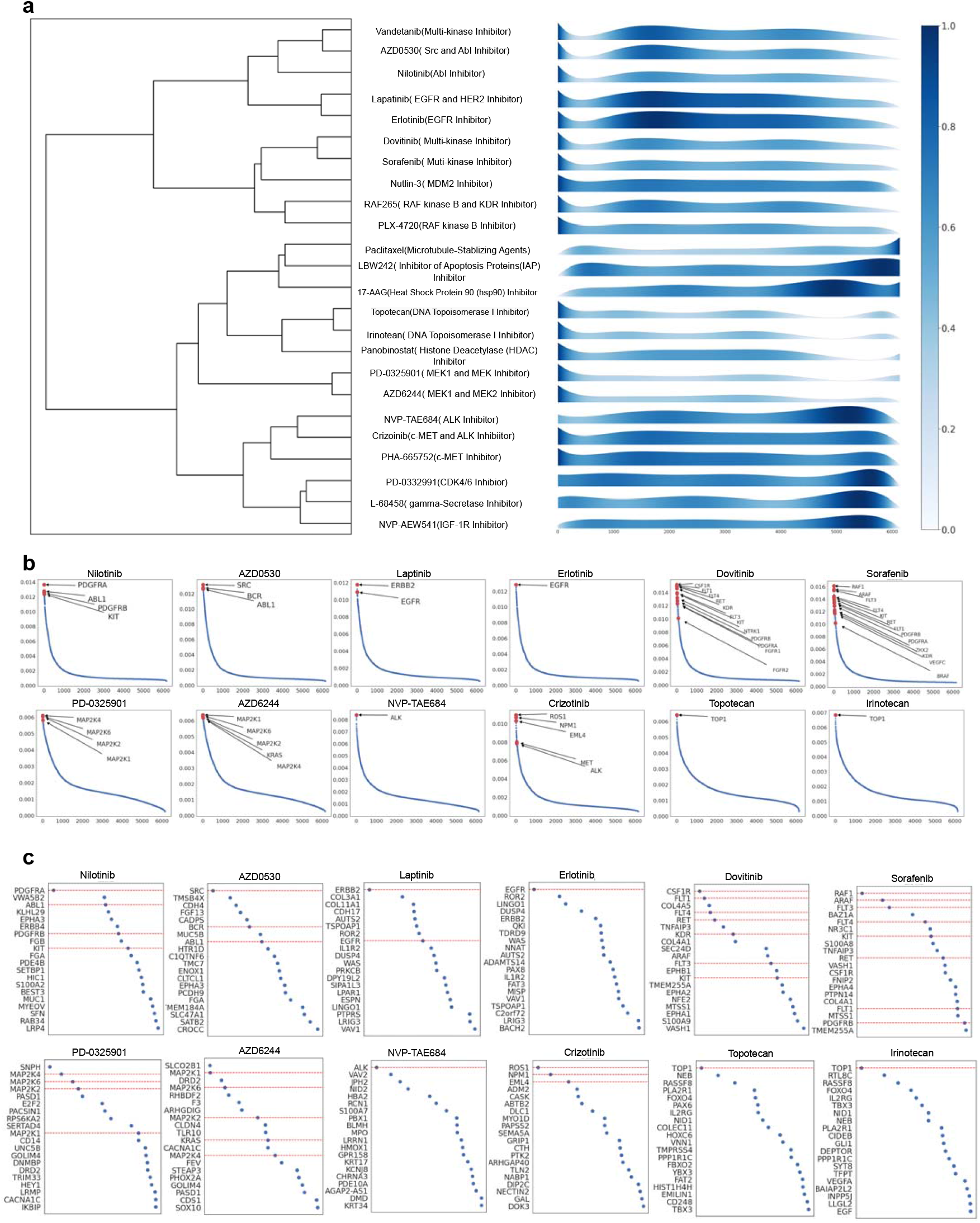
Cross Attention for Drug-Gene Targeting Mechanism Evaluation. **a**. Left: hierarchical clustering on drugs based on drug-gene cross-attention scores; Middle: the corresponding drug names and drug MOAs; Right: the drug-gene cross-attention distribution fitted by polynomial regression, x-axis: all the 6144 genes listed in a fixed order, y-axis: the fitted gene attention scores for all drugs. **b**. Gene cross-attention score for each drug in ranking order, and each drug pair is selected to have the same MOA. **c**. Top 20 genes with the highest cross-attention score for each drug.

This demonstrates the power of model learning abilities in interpreting the drug-targeting effect based on the unique cross-attention architecture design for MIDI.

### TKI Drug Target Effect Prediction on ABL&BCR Fusion Leukemia Cancer Cell Lines

CML(Chronical Myeloma Leukemia) is the most common type of Leukemia, with 30% of patients with ABL/BCR fusion mutation, this biology procedure is illustrated in Fig. 4a. Specifically, gene ALB in chromosome 9 translocating to chromosome 22 and fusing with gene BCR, resulting in a mutated chromosome called the Philadelphia Chromosome, this mutation stops the apoptosis process of normal blood cells and causes cell proliferation. TKI (Tyrosine Kinase Inhibitor) is a drug category for chemotherapy that targets the APT binding process that happens in the Leukemia cell cycles. Nilotinib is the first-generation ABL target TKI, which successfully treated 30% of Leukemia patients with ABL/BCR fusion mutation. Due to the tumor ABL drug resistance, second-generation drugs like saracatinib were developed that target SRC/ABL mutation. In this experiment, we evaluate the model’s ability to identify the TKI drug effect on ABL mutant Leukemia cancer cell lines against wild-type Leukemia cancer cell lines.

**Fig. 4:**
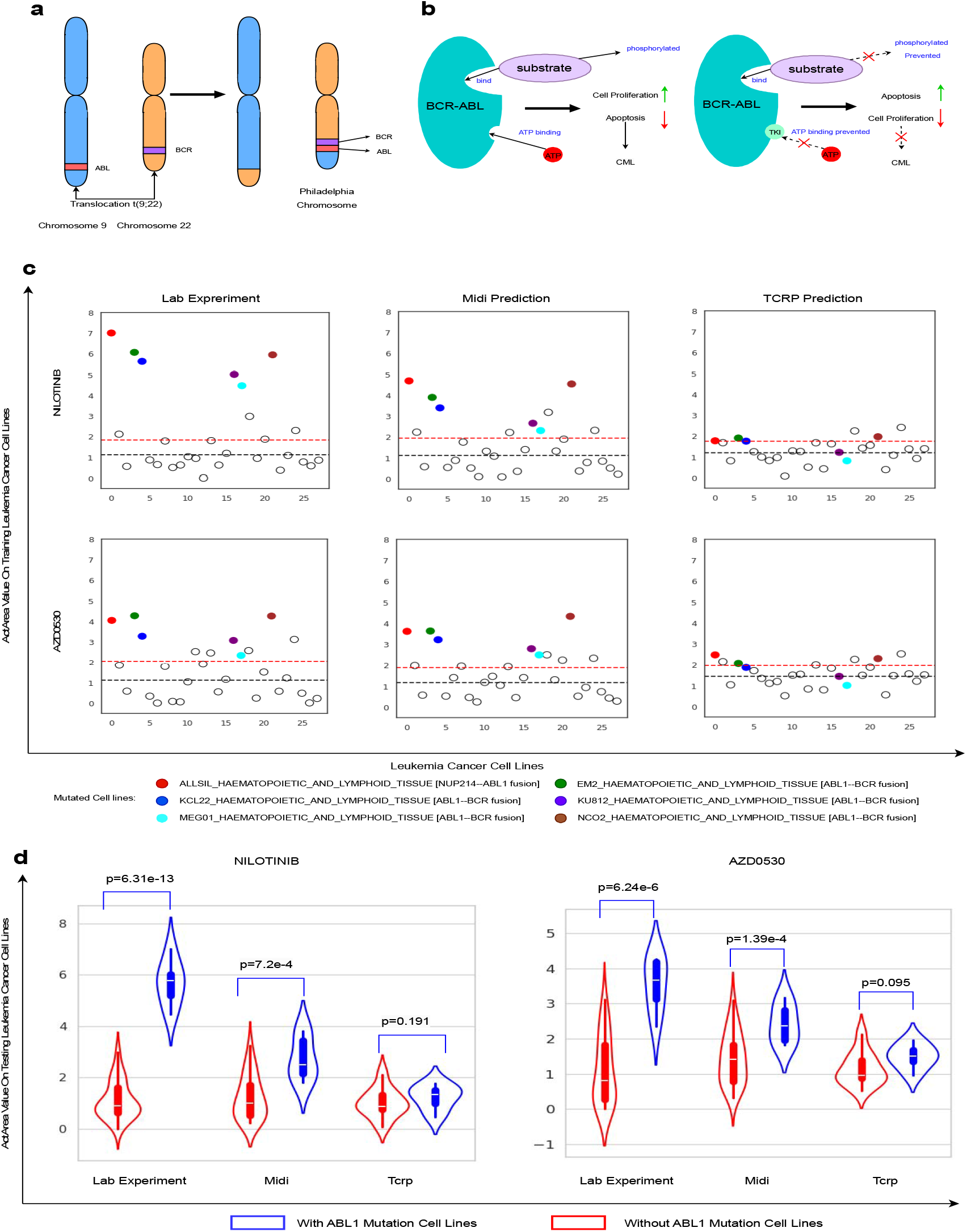
Drug Targeting Effect Prediction for Leukemia. **a**. Common Mutation Process for CML. **b**. TKI drug MOA (Mechanism of Action) process illustration. **c**. Drug effect prediction on Leukemia cancer cell lines: The x-axis is randomly ordered cancer cell lines, and the y-axis is activity area values. All the colored dot predictions correspond to ABL1 mutated cancer cell lines, and the empty black dots are predictions for wild-type cancer cell lines. The dashed black lines are the mean value for wild-type predictions, and the dashed red lines are the standard deviations for wild-type predictions. **d**. Drug effect prediction on unseen Leukemia cancer cell lines, the p-value is derived from two-sample t-tests.

We divide the experiment into two parts. For the first experiment, we use all cancer cell line data as the training data to fully evaluate the model target effect ability since it shows how well the model understands the given existing data. The training strategy is the same as in the section on cross-attention gene selection. After training, we collected all 28 Leukemia cancer cell lines in the data, within which six are ABL1 mutated. We first plot the actual drug effect from the lab experiment shown in the left column of Fig. 4b. We can see that both Nilotinib and AZD0530 show high drug inhabitation responses to all the Leukemia cancer cell lines with ABL1 mutation compared to Wild-Type cancer cell lines, this observation is consistent with the biology process of the drug MOA when they were originally designed. We then plot the drug response predictions from MIDI and TCRP, shown in Fig. 4c, in the middle and right columns. From the plot, firstly, MIDI prediction shows a clear distinction between the drug effects for both Nilotinib and AZD0530 on the ABL1 mutation cancer cell line group and the non-mutation group, all predictions are above the standard deviation of the non-mutation predictions distribution, this shows the model correctly identifies the drug effect on the targeted mutated gene. However, the predictions from TCRP don’t show a clear distinction between the mutation group and the non-mutation group, many of the prediction values fall below the standard deviation of the wild-type drug effect predictions. This shows that TCRP can’t identify the drug effect given the targeted gene mutation information, thus indicating its inability to interpret the drug-gene targeting mechanism. For the second experiment, we evaluate the generalizability of the model drug-gene targeting effect on unseen Leukemia cancer cell lines.

Specifically, we dive the data into training and testing groups. We first collect all types of cancer cell lines except Leukemia as one part of the training data, we then take 3 out of 6 ABL1 Leukemia mutated cancer cell lines and 15 out of 22 other mutated wild-type Leukemia cancer cell lines as another part of the training data, all other leukemia cancer cell lines(3 of ABL1 mutated and 7 of Non-ABL1 mutated wild-type cell lines) are collected as the testing data. Fig. 4d shows the prediction results on the testing data after model training. We perform a two-sample t-test to evaluate the model’s ability to distinguish the ABL1 mutation and the wild-type cancer cell line groups. The two groups for the actual lab experiment are clearly distinguishable with p-value=6.31e-13 for Nilotinib and p-value=6.24e-6 for AZD0530. The prediction from MIDI shows a clear distinction between the two groups, with p-value=7.2e-4 for Nilotinib and p-value=1.39e-4 for AZD0530, which aligned with the lab experiment. In contrast, TCRP can’t distinguish the two groups with p-value=0.191 for Nilotinib and p-value=0.095 for AZD0530. The testing results demonstrate the generalizability of MIDI prediction on unseen Leukemia cancer cell lines.

### Self-Attention-Based Mechanism Interpretation of Drug Molecular Binding Atoms on Targeted Genes

In this experiment, we demonstrate the model’s ability to interpret the drug MOA by identifying its major binding atoms to the protein residuals corresponding to its targeted genes. Identifying the binding mechanism is crucial for cancer drug discovery since it identifies the leading compound that functions as the fundamental chemical building block for the targeted mutated genes. We specifically pick three drugs for illustration: Nilotinib, NVP-TAE684, and Crizotinib. For the rest of the drug analyses, please refer to the supplementary materials. Fig. 5a shows the main results. Specifically, we first scan through the PDB (Protein Data Bank)^15^ by inputting the drug names and the targeted genes and find the publications with the corresponding drug structures together with the drug chemical binding mechanism on the targeted genes, we downloaded the protein-drug bonding structure file and visualize it in Pymol^16^ as shown in Fig. 5a, we highlight the binding mechanism between the atom and protein residual as orange and purple edges, which represents hydrophobic interaction and hydrogen bond respectively, the visualized protein structure is generated from one specific PDB ID listed in the table on top, we pick this specific one for illustration since it overleaps most of the common binding atoms through different protein poses listed in Fig. 5b. The plot of Fig. 5a in the top and bottom are two view perspectives of the actual 3D protein-drug binding structure, the middle plot shows the 2D drug chemical structure of highlighted top atoms with highest self-attention scores detected from the Drug Graphformer in MIDI on diagonal atom score and atom-bond scores, we pick top 5 atoms with top 5 atom bonds in this experiment. Fig. 5c shows the visualized self-attention matrix for a specific drug. For detailed top atoms picking algorithm, please refer to supplementary material. Next, to further examine the stability of the model’s ability to detect the drug molecule binding mechanism, we align the two protein structures corresponding to the PDB IDs shown in Fig. 5d for NVP-TAE684 and Crizotinib. Since these two drugs have the same gene target, the corresponding protein structures are identical, specifically the Human Anaplastic Lymphoma Kinase. Fig. 5d shows the alignment result, the rotation and translation of the two drug structures are modified accordingly to the protein alignment.

**Fig. 5:**
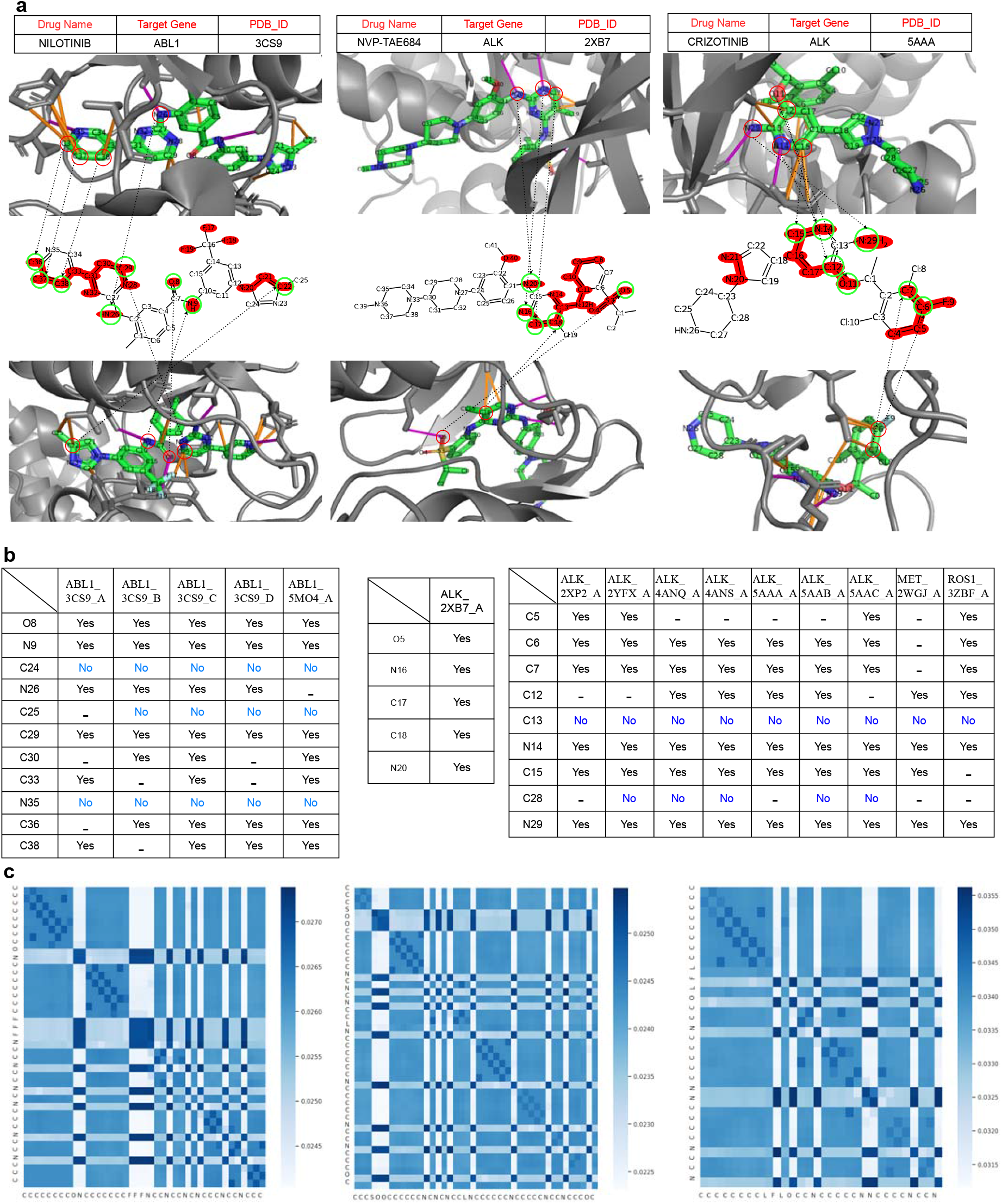
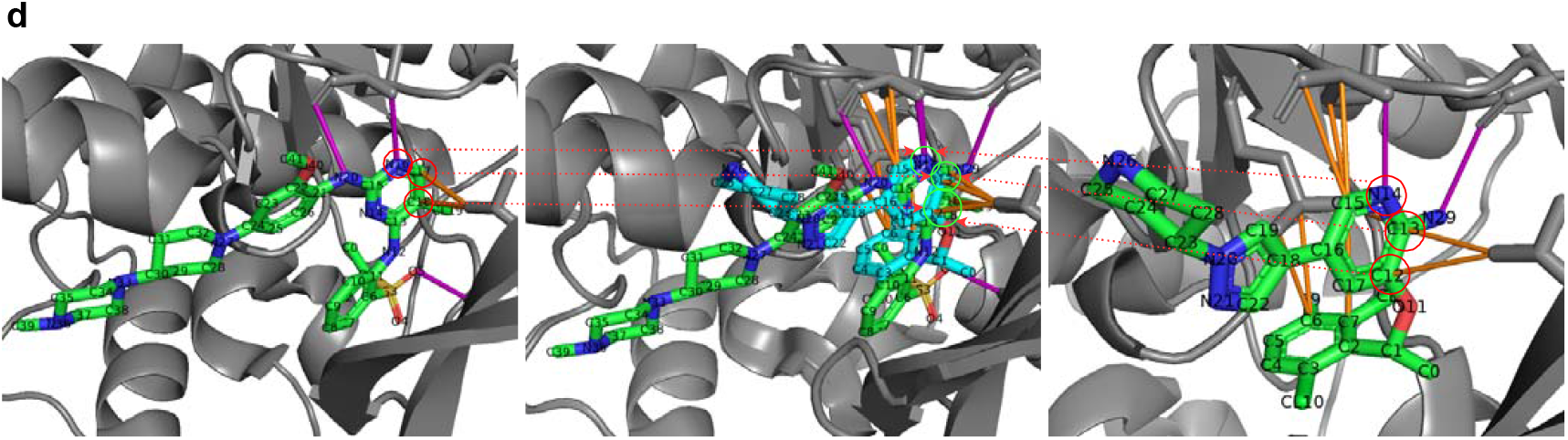
Self-Attention-Based Mechanism Interpretation of Drug Molecular Binding Cite on Target Genes. **a**. From top to bottom: Table that shows the drug name, targeted gene, and corresponding PDB_ID; One view of the protein-drug structure with atoms number and actual bindings displayed; Highlighted binding atoms selection through top atom and bond attention scores from the drug graphformer self-attention matrix; Another view of the protein-drug structure. From left to right: three different drug-protein structures. The atoms in green circles are the correctly detected binding atoms corresponding to the actual binding atoms in red circles. **b**. Tables show all the actual binding atoms (rows in the table) for drugs, the drug from left to right: Nilotinib, NVP-TAE684, and Crizotinib. Each column in the table represents one protein pose. For every entry in the table: “Yes” means the model detected that specific binding atom, and “No” means the model failed to detect that binding atom. **c**. Visualization of the self-attention matrix from MIDI for each drug. **d**. From left to right: Drug-protein structure for NVP-TAE684 and ALK; Aligned drug-protein structures from NVP-TAE684 and Crizotinib with ALK; Drug-protein structure for Crizotinib and ALK.

From plot Fig. 5a, we observe that most of the actual binding atoms are detected by MIDI by ranking self-attention atom and bond scores. Specifically, for Nilotinib, the targeted gene is ABL1, which has been discussed in Section TKI drug target effect. The binding atoms for ABL/BCR fusion mutated proteins are listed in the left column of Fig. 5b. MIDI correctly detected all the binding atoms except for C24, C25, and N35. For NVP-TAE684, whose targeted gene is ALK, MIDI detects all of the actual binding atoms shown in the middle column of Fig. 5b. For Crizotinib, whose targeted gene is also ALK, MIDI correctly detects all binding atoms except for C13. From plot Fig. 5d, after the alignment, we observe the common unique small molecule structure with the same binding atoms that exist both in NVT-TAE684 and Crizotinib, specifically N16, C17, C18 for NVT-TAE684 and N14, C12, C13 for Crizotinib, all of these atoms except for C13 are detected by MIDI. For specific statistical evaluation details, please refer to supplementary materials.

From the detection results, firstly, the atom binding detection success rate is high. For the missed-detection cases for Nilotinib and Crizotinib, even though these atoms are not exactly detected, they all belong to the same small molecule structures that contain many detected binding atoms. Specifically, for Nilotinib, N35 belongs to the ring structure containing detected atoms C36, C37, and C38; C24 and C25 belong or connect to the ring structure containing detected atoms C22. For Crizotinib, C13 belongs to the ring structure containing detected atoms C12, C15, N14, and N29. Secondly, there are some falsely detected binding atoms, such as detecting F17, F18, and F19 for Nilotinib. The reason for this could be attributed to the fact that the model hasn’t seen too many drug structures (24 drugs for training in total), so the unique and rare atom patterns from F17 to F19 are treated importantly. This indicates the necessity of incorporating a vast amount of drug training data for developing the Drug-Gene target interaction foundation model. However, this paper focuses mostly on the explanation part and the novel model architecture design for interpreting the Drug-Gene target mechanism and performing drug response prediction, so we leave the foundation model development incorporating vast training data for future development. Thirdly, since both NVP-TAE684 and Crizotinib target ALK, they share the same crucial small molecule structure (the ring structure with double Nitrogen) after the alignment, which exhibits the same binding atoms as shown in Fig. 5d, this binding mechanism is correctly detected by MIDI through picking the atoms with the highest self-attention scores. Interestingly, the irrelevant small molecule structures that don’t have the binding effect are discarded by the model, demonstrating the strong potential of applying the model for drug screening to identify the drug MOA.

### Drug Effect Prediction on General Drug-Targeted Muted Genes

For this experiment, we evaluate the drug effect prediction on types of cancer cell lines with the targeted genes mutated and compare this effect against the drugs applied to wild-type cancer cell lines. Drug effect prediction evaluation on the targeted mutated gene is important since it quantitively evaluates the model’s ability to interpret the targeting effect. Specifically, we collected 19 Drug-Target Mutated Gene pairs with the corresponding cancer types. The table in Fig. 6d shows all these pairs, note all the drugs listed in the tables had been FDA-approved, and their MOA had been well studied in targeting the specific genes (listed in the same column as the drugs) when they are mutated associated with cancer types. The last row of Fig. 4d lists the references to these targeting effects^17-34^. The specific procedure for this experiment is as follows: For each cancer type, drug, and targeted gene in the same column listed in Fig. 4d, we divide the cancer cell lines corresponding to that cancer type into two groups, the first group is the cell lines with that gene mutation, the second group is the wild-type cell lines without that gene mutation. We divide the evaluation into three groups, the first group is the ground truth lab experiment, where we take the average value of the actual drug effect applied to the gene-mutated cancer cell lines minus the average value of the actual drug effect applied to the wild-type cancer cell lines, the result is shown in Fig. 6a (left figure), we observe that all of the real lab experiment have obvious increased drug effect when it is applied to the cancer cell lines with its targeted gene mutated. We repeat this procedure with the drug effect model prediction for MIDI and TCRP. The results are shown in Fig. 4a (middle and right figures). MIDI correctly predicts all the drug effect directions when the targeted gene is mutated, which aligns with the lab experiment, while TCRP fails to predict multiple drug effect directions, specifically on the pairs: Lymphoma, NVP-TAE684, ALK; Breast Cancer, Lapatinib, EGFR; Gastric Cancer, Crizotinib, MET. More importantly, TCRP fails to predict the scale difference of drug effect between targeted gene-mutated cancer cell lines with wild-type cancer cell lines, most of its predictions on the drug effect difference are very close to zero, and only a few show moderate differences, specifically on pairs: Leukemia-Nilotinib-ABL1 with 0.8; Leukemia-AZD0530-ABL1 with 0.8; Leukemia-AZD0530-SRC with 0.5. However, the scale of the predicted drug effect differences is still very small compared to the lab experiments, specifically 4.8, 2.8, and 2.7 respectively. Fig. 6b

**Fig. 6:**
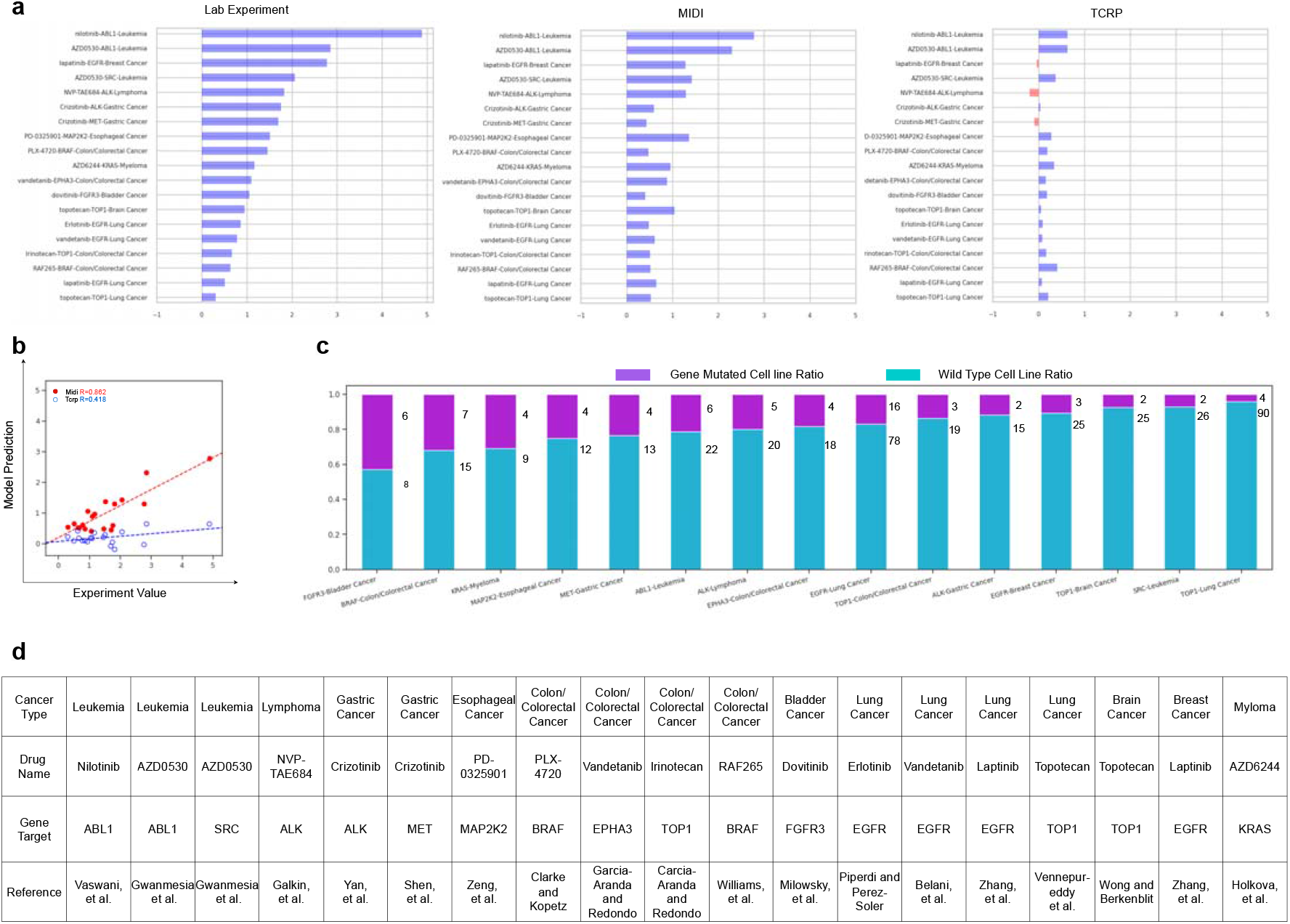
Cancer Drug Effect Prediction on Targeted Mutated Genes. **a**. Drug target effect prediction performance comparison, the average drug effect prediction difference between mutated and wild-type cancer cell lines. The x-axis: activity area value difference. The y-axis: all the drug-gene target effect pairs on specific cancer types. **b**. Scatter plot of correlation comparison for the drug effect prediction difference between mutated and wild-type cancer cell lines. **c**. The ratio between gene mutated and wild-type cancer cell lines. The number beside the ratio is the actual cancer cell line number. **d**. Documented references that the actual lab experiments had been performed to validate the drug MOA and its target genes.

## Discussion

The massive improvement of MIDI on per-drug effect prediction accounts for its unique architecture design. Specifically, the unique cross-attention feature selection mechanism introduced in Section Cross-Attention Design. This mechanism enlarges the effect of certain targeted gene embeddings interacting with the drug embeddings, which acts as a guided feature selection procedure^35^, so the self-attention mechanism for the drug puts more attention on those specific targeted genes’ information (gene expression values, mutation information, and identification embeddings). It makes changes to the model parameters accordingly in the learning phase to recognize the unique drug structure by itself on identifying different targeted genes. Hence, the model is more sensitive to the heterogeneities of the individual Drug-Cancer Cell Line pairs by focusing on the unique gene targets. More specifically, the drug graphformer self-attention model parameters are updated accordingly and put more weight on important molecular structures and atom embedding so that they are much closer to the targeted gene embeddings in the latent space and counted for major changes to the drug effects. We excitingly find this self-attention mechanism can identify the important binding atoms that bind the specific protein residual of the targeted genes corresponding to a drug structure, which is specifically discussion in later section.

## Method

### Model Inputs

Fig. 1A shows the details of the model architecture, the use case for interpreting drug MOA, gene target ranking, and the drug effect prediction process. The input for MIDI contains two parts: the canonical drug SMILE sequence and the bulk cancer cell line information. We use the gene information associated with each cell line as the second part of the MIDI input, including gene names, gene expression values, and mutation information. The two parts of the inputs are separately discussed in the following sections.

### Drug Inputs

To represent the drug information, we use the canonical SMILE sequence as the drug input^36^. We apply Drug Graphformer Architecture^37,38^ to process the drug SMILE information. To generate proper input for the Drug Graphformer, we made further processes on the SMILE sequence. We first use the Rdkit package^39^ on the SMILE sequence to obtain the bonding information between atoms and their edge type information, the bonding information is obtained as a connectivity adjacency matrix, with the edge type associated with it. As shown in Fig. 1A and Fig. 1C, the inputs from the SMILE sequence to the Drug Graphformer contain three parts specifically discussed in the following paragraphs.

The first part is the drug atoms input, we first tokenize all the atom’s names (such as “C” and “N”) into a TensorFlow vocabulary^40^, then encode these names as a *K*_*a*_-dimensional one-hot encoding vector, mathematically represent it as 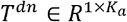, where *K*_*a*_ the dimension of the atom vocabulary. The second part is the atoms’ bonding information, represented by edge type obtained from the smile sequence. Specifically, we collect 5 edge types^41^: single bond, double bond, triple bond, aromatic bond, and no bond. These edge types are encoded into a 5-dimensional one-hot vector 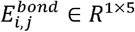. The third part is the relative position distance matrix, the structure information input for the Drug Graphformer. Specifically, given a SMILE sequence, we first obtain the adjacency matrix 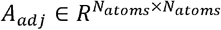 (Where *N*_*atoms*_ is the number of atoms in the smile sequence) between atoms from the Rdkit package^39^, from this adjacency matrix, we can calculate the shortest path distance between every two atoms using Dijkstra’s algorithm^42^, we treat the shortest path distance as the relative distance between two atoms. We represent the relative position distance matrix mathematically as 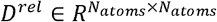. The value of *i, j* entry 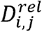 represents the shortest path distance between atom *i* and atom *j*.

### Gene Inputs

The second part of the input is the gene information corresponding to each cancer cell line, it contains three parts: gene names, gene expression values, and gene mutation information. The gene input is pre-processed to filter out those genes with zero gene expression values and low gene expression value variance.

For encoding gene name information, we directly pass the gene names into the Geneformer pre-trained model^43^ and get the embeddings as the input gene name representation. As stated in the original paper, Geneformer is a state-of-the-art pre-trained gene transformer model trained on 30 million single cells, its gene name embeddings encode the gene functionalities. The gene name embedding is represented as 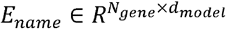, where *N*_*gene*_ is the total number of genes after pre-processing and *d*_*model*_ is the embedding dimension. For the gene expression input, it’s a single continuous value 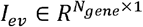 represents the expression scale of a single gene after pre-processing. *l*_*ev*_ is directly passed to a position-wise feed-forward neural network *H*_*ev*_ as *E*_*ev*_ *= H*_*ev*_ (*l*_*ev*_), where 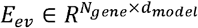 is the embedding vector for each gene expression. The gene mutation input is a two-dimensional one-hot vector representing binary information on whether one specific gene is mutated or not *I*_*mt*_ ∈ {0, 1}. We pass *I*_*mt*_ to another point-wise feed-forward neural network *H*_*mt*_ as *E*_*mt =*_ *H*_*mt*_ (*I*_*mt*_), where 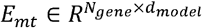 is the embedding vector for each gene mutation. The final gene embedding vector is the summation of the above three embedding vectors in the latent space shown in equation (1).

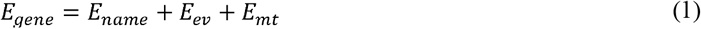

### Drug Graphformer Design

The Drug Graphformer model architecture for MIDI is shown in Fig. 1C. It is designed based on the latest approach^37^, which is mainly used for performing self-attention and generating drug embeddings. The general Graphformer message passing update function is applied as atom nodes embedding shown in equation (2).

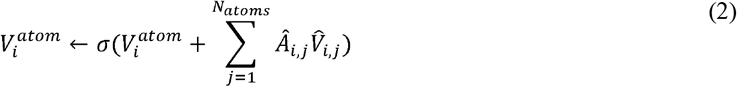

Where σ is the non-linear activation function Relu^44^, 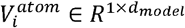 is the node embedding for the atoms, *i* is the atom index, *j* is the neighboring atom index to 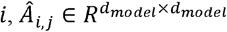 is the mixed attention matrix that incorporates three modality information between atom *i* and atom *j*, and 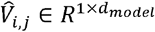 is the corresponding mixed embedding values. The attention matrix and the corresponding mixed embedding value are computed shown in equation (3) and equation (4).

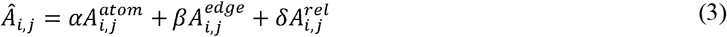

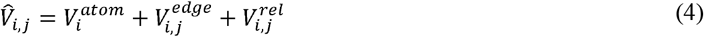

Where 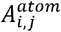 is the atom-atom attention score, 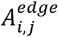 and 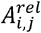 is the attention score of the *i*th atom embedding to the edge type embedding and relative position distance embedding between atom *i* and *j* respectively. 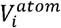 is the *i*th atom embedding, 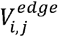 and 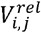 are the edge type embedding and relative distance embedding between *i*the atom and *j*th atom. *α, β* and *δ* are weighted parameters that can be customized. These mixed modalities are separately discussed as follows.

#### Atom-Atom Attention Score

For atom embedding, firstly the th atom input 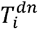 is passed through a position-wise feed-forward neural network 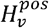 as 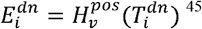, where 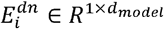. The atom-atom attention score is computed shown in equation (5).

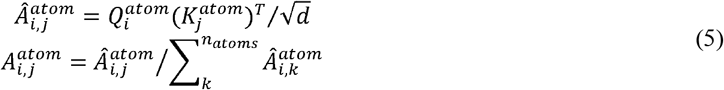

Where 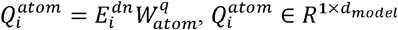 and 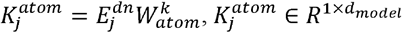 are the query and key embeddings for atom respectively, is a constant scaling factor, 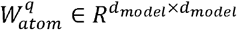, and 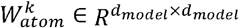. The atom embedding is computed as 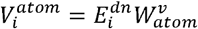.

#### Atom-Edge Attention Score

Given the edge type input 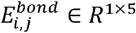, it is passed to a fully connected layer to get the edge type embeddings 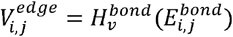, and 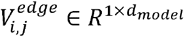. The atom-edge attention score is computed in equation (6)

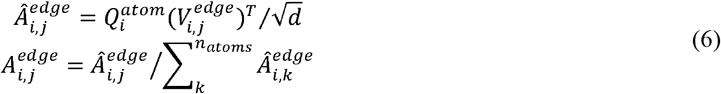

#### Atom-Position Attention Score

For the relative position embedding, we apply the same position encoding dictionary lookup scenario^45^ as shown in equation (7)

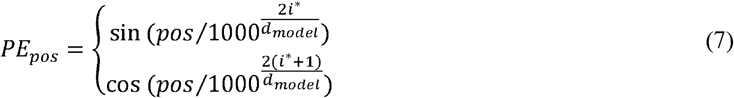

Where *i*^*^ is the dimension in latent embedding space. When computing the relative distance embedding 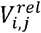 between atom *i*, and atom *j*, we take 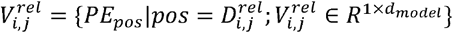, where 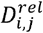 is the shortest path distance between atom and atom. The atom-position attention score is computed in equation (8)

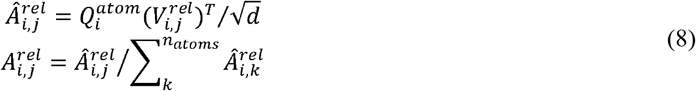

### Cross Attention Design for Drug Targeted Gene Ranking

The target gene ranking and selection for MIDI is based on sequential feature attention score^35^. Firstly, the atom embeddings are flattened to form a global embedding representation of a specific drug:

*E*_*drug*_ = *flatten* (*V*_*atom*_), where 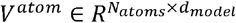 is the atom embedding representations for this drug. Because the drug mechanism of action is fixed when its chemical structure is fixed, its gene targets are fixed too regardless of the gene expression values or the gene mutation information. Therefore, we apply the cross attention between the global drug embedding with only the gene identification embeddings as shown in equation (11), Fig. 1D visually shows this process as well.

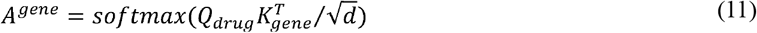

Where 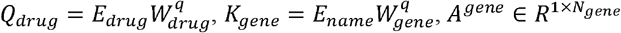, is the cross attention scores, which are used as the gene importance ranking score regarding to a drug, the softmax function in equation (11) is modified with a temperature parameter *T*^46^, which can be customized to sharp the gene distribution and put much higher attention score on the drug targeted genes, we specifically set *T* = 9. Fig. 3B visually shows the sharp gene ranking distribution with this modification. We demonstrate that with the unique cross attention design between the drug chemical structure embedding and the gene identification embedding, MIDI could incorporate the prior drug-gene target knowledge to help identify the drug binding mechanisms, classify different drug types, and possess the potential to identify gene target regarding to a specific unknown drug chemical structure based on a data-driven training scenario.

After obtaining the cross-attention score, *A*^*gene*^ is directly multiplied to each gene embedding *E*_*gene*_ to get the scaled gene embedding representation shown in equation (12).

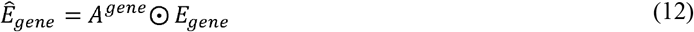

where ⊙ represents element-wise multiplication, *A*^*gene*^ functions as the coefficient score on selecting important genes, which is demonstrated to be equivalent to the feature selection in Sequential Lasso^35^.

### Adding Prior Drug-Gene Interaction Knowledge

#### Model Prediction and Loss Function

As shown in Fig, 1A, the output of prediction of MIDI is a single numerical value representing the drug effect applied on a certain cancer cell line, specifically the Activity Area^47^, we compute this value by combining both the drug information and the gene information within the cancer cell line. The global drug embedding *E*_*drug*_ and the scaled gene embeddings *Ê*_*drug*_ are separately passed through feed-forward projection layers *Pr*_*drug*_ and *Pr*_*gene*_ to get two single numerical values shown in equation (13)

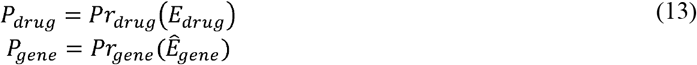

Where *P*_*drug*,_ *P*_*gene*_ ∈ *R*1, These two values are added together to form the final Activity Area prediction value *P*_*de*_ as the drug effect shown in equation (14)

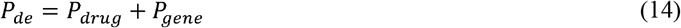

The loss function contains three parts as shown in Fig. 1B. Firstly, Mean Square Error Loss is used to approximate the actual Activity Area value in the training data as shown in equation (15).

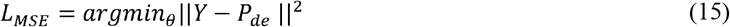

Where *Y* is the true Activity Area value, *θ* represents all model parameters.

Secondly, Supervised Contrastive Loss is applied to incorporate the prior drug-gene interaction information as shown in equation (16).

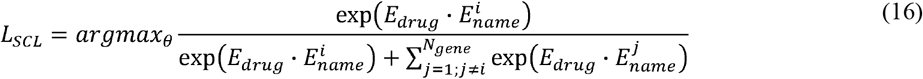

Where *i* is the gene index corresponding to the genes targeted of *E*_*drug*_, *j* are all the non-target gene index. The maximization of *L*_*SCL*_ ensures the embedding of a certain drug much closer to its targeted gene identification embeddings than other non-targeted gene embeddings in the latent space. Therefore, when performing the cross-attention gene ranking introduced in equation (11), the model could put more attention weights and focus more on those targeted genes.

Thirdly, we adopt a Self-Supervised Contrastive Loss as a regularizer for distinguishing drug embeddings in the latent space shown in equation (17).

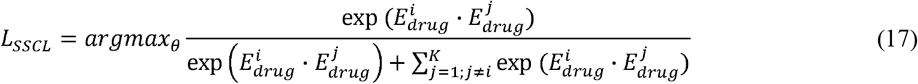

Where *K* is the drug input batch size, *i* is the index of a specific drug in one training batch, *j* is the index of all other drugs different to drug *i*. Maximizing *L*_*SSCL*_ ensures the distinguishment in the embedding space of different drug structures. Combining *L*_*SSCL*_ and *L*_*SCL*_, when different drug structures target same gene sets, the model could firstly enforce the recognitions of those targeted genes associated with the drugs, meanwhile remaining unique distinguishment of different drug chemical structures in the embedding space.

Fig. 1B visually shows the details of the above two loss as well, the final loss function is the combination of the above three losses as shown in equation (18).

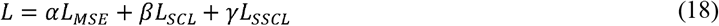

Where *α, β, γ* are customized parameters that weights the importance of each loss, we specifically set *α* = 1, *β* = 1, *γ* = 0.2 to put more focus on prediction accuracy and incorporating prior knowledge.

